# Subjective inflation of color saturation in the periphery under temporal overload

**DOI:** 10.1101/227074

**Authors:** Sivananda Rajananda, Megan A.K. Peters, Hakwan Lau, Brian Odegaard

**Author notes:** **Correspondence should be addressed to:** Sivananda Rajananda,UCLA Psychology Department,1285 Franz Hall,Los Angeles, CA 90095-1563.

## Abstract

A long-standing puzzle in perception concerns the subjective impression of vivid color experience in the periphery. While peripheral color processing is not entirely insensitive, the subjective vividness seems unsupported by the relative paucity of representation in the periphery. Inspired by the flashed face distortion effect, in which peripheral faces are perceived as somewhat exaggerated or distorted when they are preceded by flashes of other faces in the same location, we attempted to create an equivalent illusion in the domain of color. The hypothesis is that under temporal overload, patches of colorful dots may be perceived as more exaggerated in terms of saturation. We confirmed this hypothesis with the observation of a significant effect of modest magnitude, which was replicated in a second experiment. These results suggest that subjective inflation of perceived color saturation does occur in the periphery, when the perceptual system is sufficiently occupied temporally and spatially. We discuss the relationship between the observed effects with previous findings of liberal detection biases in the unattended periphery.

---

Do we perceive the visual periphery in precise detail, or do we overestimate the resolution with which we perceive the visual surround? While some researchers claim that we perceive fine details in the periphery (Haun, Tononi, Koch, & Tsuchiya, 2017; Kaunitz, Rowe, & Tsuchiya, 2016; Vandenbroucke et al., 2014), others disagree (Cohen, Dennett, & Kanwisher, 2016; Dehaene, Changeux, Naccache, Sackur, & Sergent, 2006; Kouider, de Gardelle, Sackur, & Dupoux, 2010). This tension in the literature has led some of us to hypothesize that the subjective phenomenology of peripheral perception may be ‘inflated’, in the sense that we experience or think we see more than we actually perceptually represent (Lau & Rosenthal, 2011).

One line of evidence in support of this peripheral inflation account is that at equivalent sensitivity, subjects use a more liberal criterion for detection in the periphery than in the center (Solovey, Graney, & Lau, 2015). That is, subjects say they see the target (e.g., a gabor patch) more often in the periphery, even though they are not better at detecting it in terms of sensitivity or the capacity to produce correct responses. A similar effect has been observed when we compared detection at endogenously cued vs uncued locations (at equivalent eccentricity), and has been replicated several times under different conditions (Rahnev et al., 2011).

It has been argued that detection biases can reflect subjective aspects of perception (Peters, Ro, & Lau, 2016; Witt, Taylor, Sugovic, & Wixted, 2015), and these results (Rahnev et al., 2011; Solovey et al., 2015) have been interpreted as such. In particular, these effects are unlikely to reflect strategical cognitive or reporting biases, because they persisted even after hundreds of trials of training with behavioral feedback. However, there is still a sense that the connection to subjective phenomenology may be indirect. If liberal detection biases in the periphery truly reflect subjective inflation at the phenomenological level, one should be able to create more straightforward visual illusions based on this underlying mechanism, outside of the context of detection tasks near psychophysical thresholds.

One illusion that may be related to these effects is the flashed face distortion effect (Tangen, Murphy, & Thompson, 2011): When eye-aligned faces are flashed continuously in the peripheral visual field, the faces start to look like caricatures of themselves, whereby different features of the face appear subjectively to be highly exaggerated and even grotesque. This phenomenologically vivid illusion may be related to the detection bias effects mentioned above because detection requires recognizing a stimulus as being different from the background or a norm. If a person’s facial features are detected with more liberal biases, such features may also look more exaggerated phenomenologically, leading to the grotesque impressions.

We reasoned that if this analogy is true, we should be able to find the equivalence of the flashed face distortion effect in other domains (i.e., with other stimuli) where subjective inflation generally occurs. Here, we have chosen to focus on color saturation. It is known that at identical stimulus size, color sensitivity is poorer in the periphery than in the central visual field (Hansen, Pracejus, & Gegenfurtner, 2009). And yet, we do not tend to subjectively see peripheral stimuli as having lower saturation (i.e. as less colorful), suggesting that peripheral inflation is probably at play. Therefore, here we tested whether we might be able to find an equivalent illusion in color space when using a setup similar to the flashed face distortion effect.

## Experiment 1

One critical feature of the flashed face distortion effect (Tangen et al., 2011) is that the illusion occurs only after a few repetitive flashes of different faces in the same locations in the periphery. Here we tested if this kind of temporal overloading would lead to color patches being perceived as more deviant from the norm or background, too (i.e., whether they would look more vividly colorful compared to a standard of the same saturation at a central location, which would not be preceded by flashes of similar color patch stimuli).

### Methods

#### Participants

Eighteen participants were recruited online through Amazon Mechanical Turk and participated in Experiment 1. Participants who completed the task were compensated with $2 each with an incentive of an extra $1 if they performed better than the previous participant. Participants who did not fully complete the task were paid a prorated amount to compensate them for their time. All participants were required to read an online consent form before participating, and verify that they did not have a history of any neurological conditions (seizures, epilepsy, stroke), had normal or corrected-to-normal vision and hearing, and had not experienced head trauma before participating in the experiment. The University of California Los Angeles IRB granted ethical approval for the study; all procedures were performed in accordance with the Declaration of Helsinki.

#### Stimuli

Each stimulus was composed of an image with a gray background and 3 patches of colored dots (with dimensions of 150 x 150 pixels per patch) aligned along the horizontal midline of the screen. The middle patch was located in the center of the screen, and the side patches were located approximately 10 degrees to the left and right of the vertical midline. To approximate the location of the patches in visual degrees, each participant was prompted to estimate the distance between their eyes and the monitor in inches, and enter this information before the start of the experiment; from this estimate, the distance between the patches in pixels was calculated to correspond to the distance in visual degrees. Each patch was composed of 30 distinct dots (see Fig. 1), with each dot having a 14-pixel radius. The coordinates of the center of each dot were chosen randomly within each patch, and the radius of each dot was only allowed to overlap by a maximum of 30% (4.2 pixels) with neighboring dots, to simultaneously minimize white space within a patch while maintaining visibility of each individual dot. Following the HSV (Hue, Saturation, Value) model of color, the value of all dots was set to 1.0, and the hue was randomly drawn for each of the dots within each stimulus. The saturation of the dots in the center patch was 0.5, while the saturation of the dots in the side patches were either 0.3, 0.4, 0.5, 0.6, or 0.7, depending on the trial. All of the dots in both the left and right side patches had the same saturation within each stimulus on a given trial.

**Figure 1.**
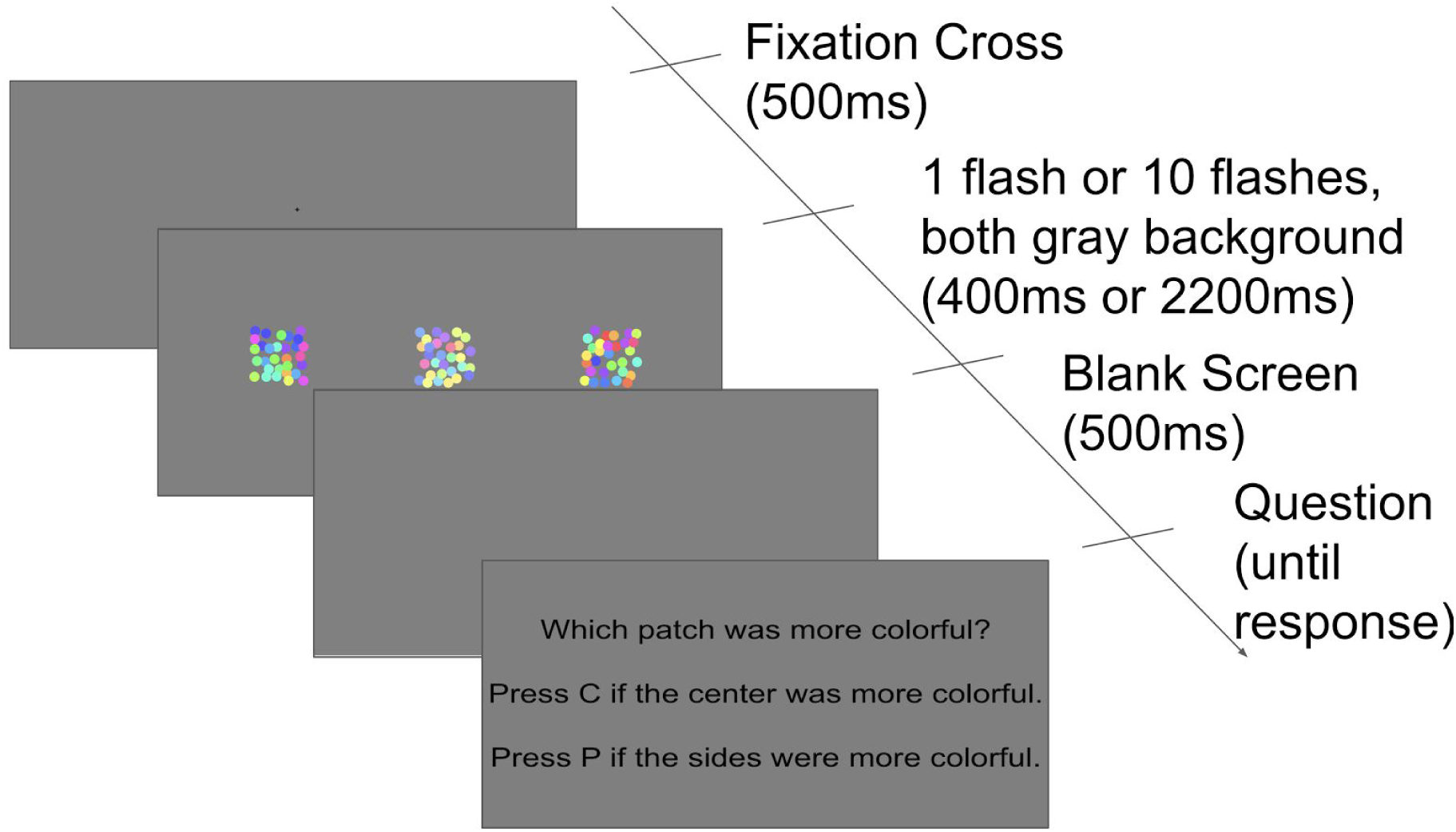
Stimulus presentation timeline for Experiment 1. Each trial began with the display of a fixation cross at the center of the screen. After 500ms, subjects were presented with either one flash of the three patches, or 10 sequential (0ms ISI) presentations of the patches in the periphery, with the dots in the peripheral patches changing in both hue and position (but not saturation) on each flash. In the 10-flash condition, the center patch was constant throughout the peripheral flashing. Following a blank screen, the trial concluded with the question prompt, asking subjects to indicate which was more colorful: the last peripheral patches, or the central patch.

#### Procedure

Before the experiment began, illustrated instructions were presented to ensure that the participants understood what the word ‘saturated’ meant, and could view examples of the trial stimuli. During the experiment, each trial began by displaying a fixation cross in the center of the screen for 500ms (see Fig. 1). Subjects were instructed to keep their eyes on the cross in the center of the screen throughout the duration of each trial. Following fixation, the stimulus (i.e. images consisting of the three patches) was flashed once for 400ms (“1-flash Condition”) or 10 times for 200ms each with 0ms ISI (no blanks between images) except the last (10th) flash, which remained on the screen for 400ms (“10-flash Condition”). The rationale for a longer final flash was to provide a unique timing for the last presentation, so participants were aware that they were viewing the final example in the sequence, to compare the saturation of patches. Within each trial, the center patch remained identical across all flashes within a trial (i.e., essentially remained a constant stimulus), while the side patches were generated randomly for each flash; that is, during each flash, the position and hues of the dots on the side patches were randomly regenerated. Following the presentation of the patch, a blank screen was shown for 500ms, and then a question prompt appeared, asking subjects to identify which was more colorful (saturated): the peripheral patches (on the left and right), or the patch in the center. The subject would then key in their response on the keyboard (“p” if the side patches was perceived as more saturated, and “c” if the center patch seemed more saturated). All trials from both the 1-flash and 10-flash conditions were interleaved randomly. Each participant completed 400 trials (2 conditions × 5 saturation levels × 40 trials per saturation level). Two one-minute breaks were given: one after trial 133 and and another after trial 267.

### Results

**Figure 2.**
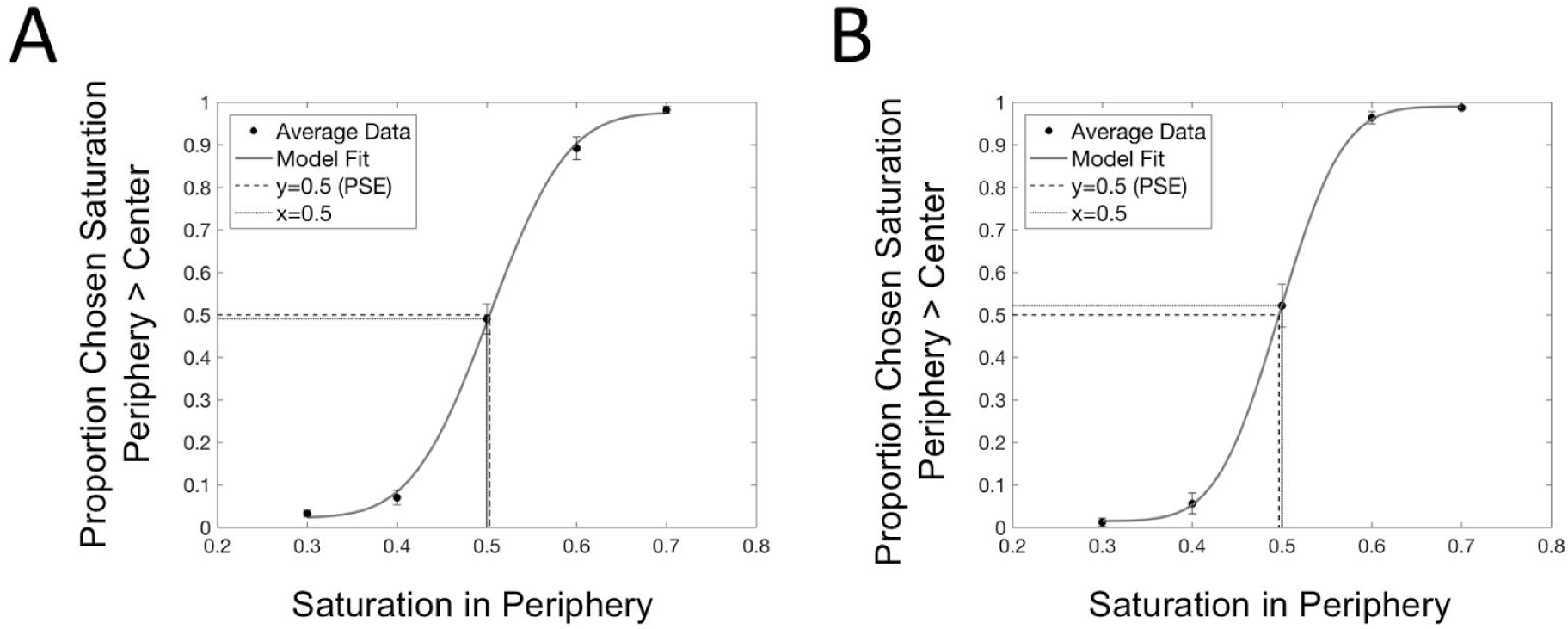
Psychometric fit to data averages across subjects. (A) Averaged data and psychometric fit for the 1-flash condition in Experiment 1, where we compared the perceived saturation of patches in the center versus periphery without preceding flashing stimuli. Error bars represent standard error of the mean (SEM). (B) Averaged data and psychometric fit to averaged data for the 10-flash condition, where the peripheral stimuli to be judged were preceded by 9-flashes before they were compared with the central standard. A two-tailed, within-subjects paired-sample t-test on the Points of Subjective Equality (PSEs) showed that subjects perceived the peripheral stimuli as being more saturated during the 10-flash condition compared to the 1-flash condition (t(15) = 2.4978, p = 0.0246). The numerical magnitude of the effect itself was however modest (1-flash PSE mean: 0.5052; 10-flash PSE mean: .4949).

**Figure 3.**
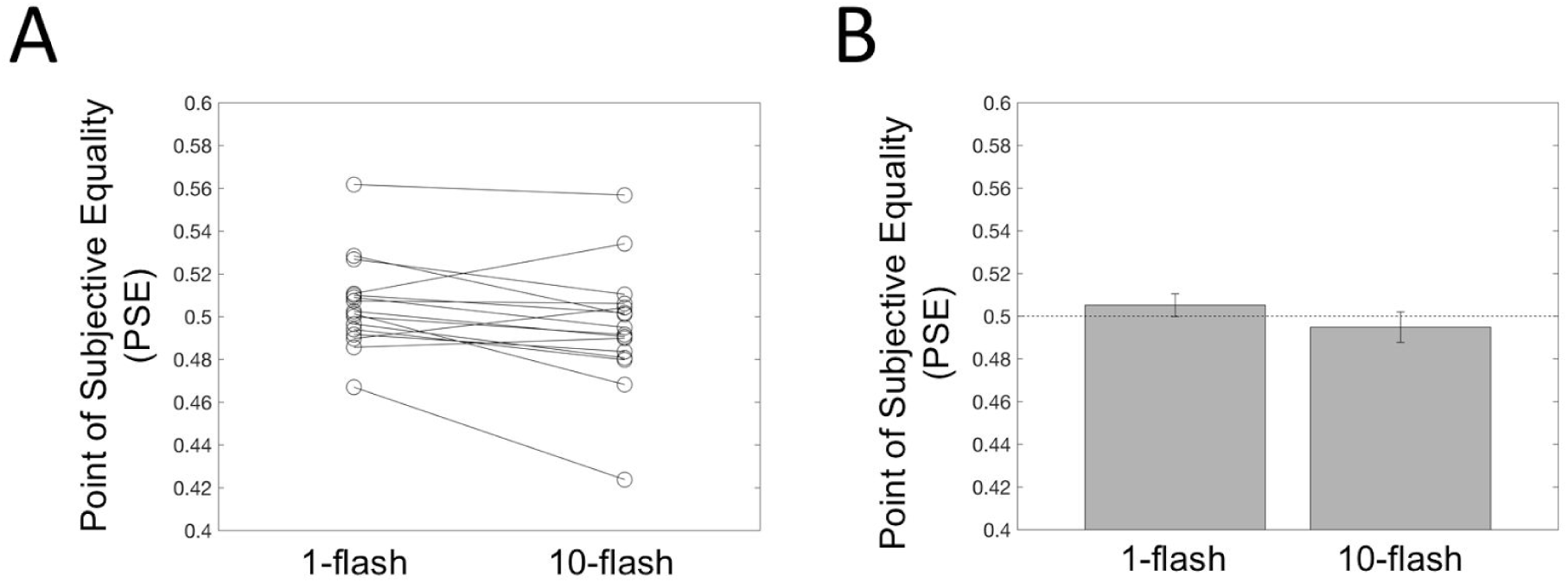
Estimates of the Point of Subjective Equality (PSE) for the 1-flash and 10-flash conditions in Experiment 1. (A) A plot of PSEs for each individual subject for the 1-flash condition and 10-flash condition. (B) Average PSEs for the two conditions. Bars display the mean PSE, and error bars represent the standard error of the mean of PSEs across all subjects.

We fitted a psychometric curve (cumulative Gaussian function) to each individual subject’s data in each of two conditions, which produced two unique psychometric fits. If at least one of the two fitted psychometric curves of a participant had a range of < 0.2 (with range defined as ((y when x=0.7) - (y when x=0.3)) or slope of < 0.1, then the data from both conditions for that participant was discarded. Based on this criterion, we discarded two subjects’ data, yielding a total of 16 subjects for our analyses. For these remaining subjects, we used the psychometric fits to calculate the Point of Subjective Equality (PSE) in each condition for each subject.

Results from a two-tailed, within-subjects paired-sample t-test on the PSEs showed that subjects perceived the peripheral stimuli as being more saturated during the 10-flash condition compared to the 1-flash condition (t(15) = 2.4978, p = 0.0246). While the magnitude of the effect was small (1-flash PSE mean: 0.5052; 10-flash PSE mean: 0.4949) and perhaps subjectively not as striking as the flashed-face distortion effect, this finding still provides preliminary support for our hypothesis: Even a feature dimension such as color can reveal distortions/inflation upon repeated presentation in the visual periphery.

## Experiment 2

We wondered if the statistically significant effect in Experiment 1 is small in magnitude because the stimuli were presented on grey background, which was not close enough to natural perception in everyday settings. Perhaps the effect of temporal overloading induced by flashes will be stronger if the visual system is also sufficiently overloaded spatially, as in naturalistic settings. Therefore, we next sought to replicate Experiment 1 with stimuli presented on a real-world image background rather than a simple gray background. We hypothesized that this manipulation would increase the magnitude of the effect found in Experiment 1; that is, participants would perceive the repeatedly-presented colors in the peripheral locations as being even *more saturated,* compared to what was seen in the first experiment, which would be reflected in a larger shift in PSE and larger effect size.

### Methods

#### Participants

Twenty-three participants were recruited online through Amazon Mechanical Turk. The payment protocol, consent procedure, exclusion criterion, and IRB approval procedures were identical to Experiment 1.

#### Stimuli & Procedure

Color patch stimuli were identical to those used in Experiment 1, with two exceptions. First, the assumed resolution of each participant’s screen was assumed to be 120 DPI (dots/pixels per inch). Second, the background changed as a function of condition. During presentation of the colored dot stimuli, the background was either a photo of a real-world scene (“Real-World” Condition) or a gray background (“Gray” Condition). The real-world scenes included pictures of a hotel room, a train station, a street in a residential area, a chapel, a kitchen with dining table, and a store window. Each participant was randomly assigned to one background photo for all trials in the Real-World condition, in order to ensure fast presentation times in their web browsers.

Each trial followed the same procedure as the 10-flash condition in Experiment 1, with all trials from both the Real-World and Gray conditions interleaved randomly. Each participant completed 200 trials (2 conditions x 5 saturation levels x 20 trials per saturation level). Two one-minute breaks were given: one after trial 67 and and another after trial 133.

**Figure 4.**
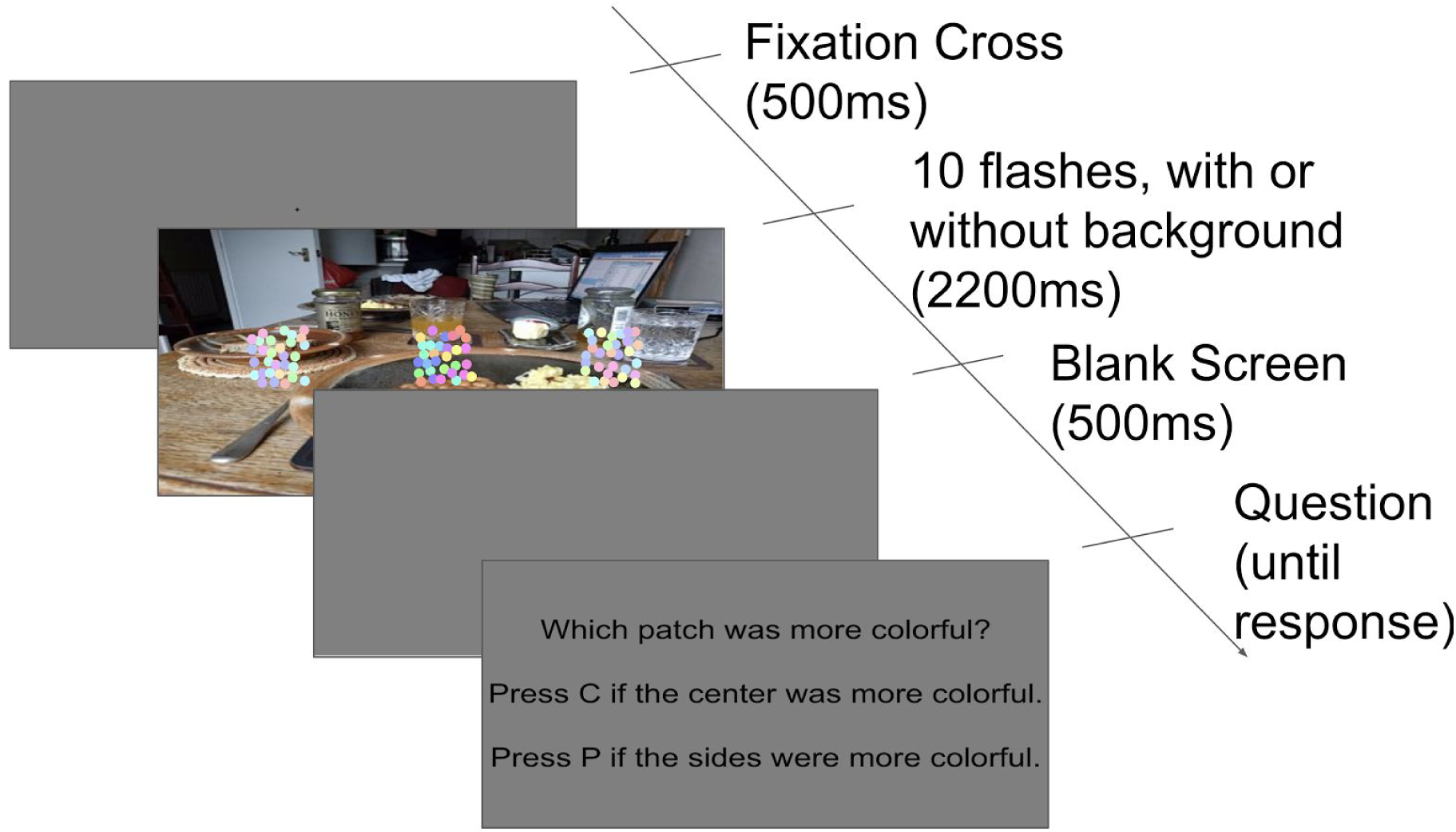
Stimulus presentation timeline in Experiment 2. The timeline in Experiment 2 was similar to Experiment 1, but in this experiment one of the conditions included the presentation of the circular dot patches on an image of a real-world scene. In both cases, the peripheral stimuli were preceded by 9 flashes (i.e., identical in timing to the 10-flash condition from Experiment 1). Shown here is a sample trial from the Real-World condition.

#### Results

**Figure 5.**
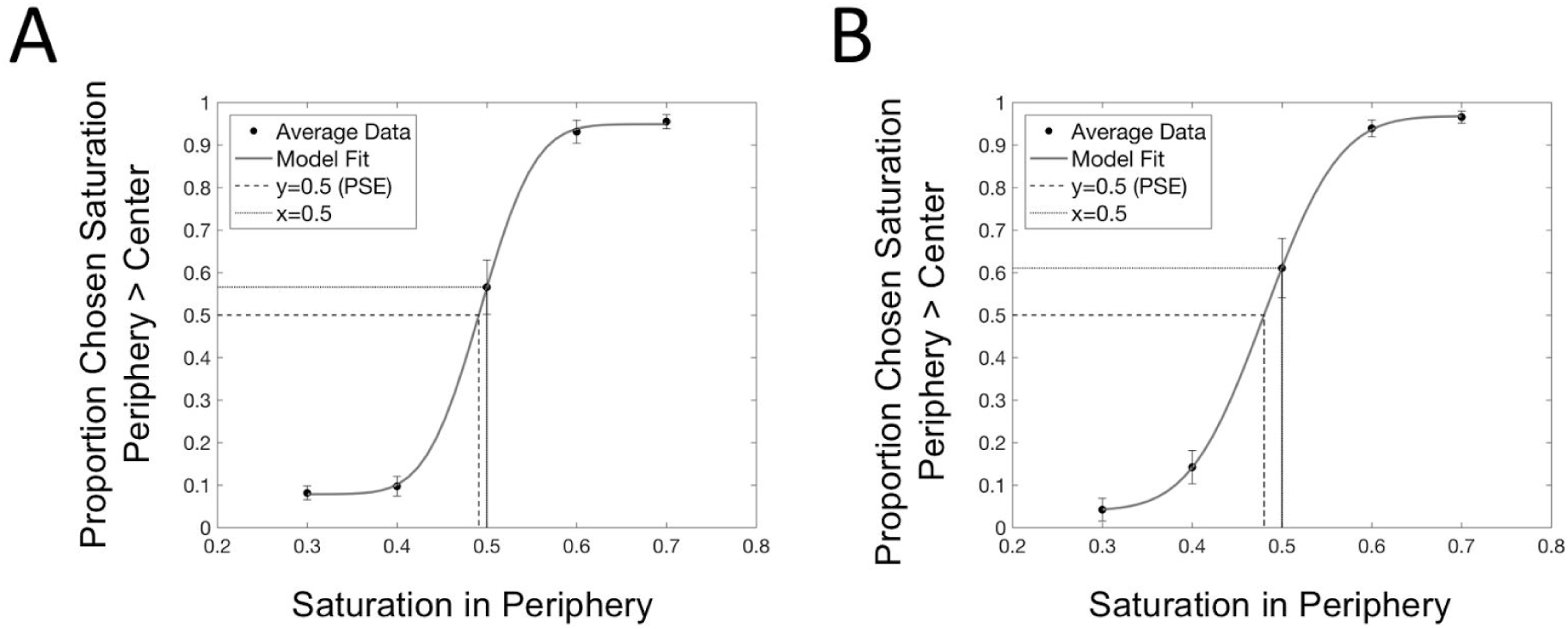
Psychometric fit to data averages across subjects in Experiment 2. (A) Averaged data and psychometric fit for the Gray condition, where the background was a simple uniform background. Error bars represent standard error of the mean (SEM). (B) Averaged data and psychometric fit to averaged data for the Real-World condition, in which the background was a naturalistic everyday image. A two-tailed, paired-sample, within-subjects t-test on the Points of Subjective Equality (PSEs) indicated that subjects perceived the peripheral stimuli similarly during the Gray condition compared to the Real-World condition similarly (t(18) =1.1643, p = 0.2595). We also conducted a 1-sample t-test of the PSE in each condition against 0.5 and found that the PSEs in the Gray condition (PSE mean: .4911) were not significantly different from 0.5, although there was a trend in the hypothesized direction (t(18) = −1.8703, p = 0.0778).

**Figure 6.**
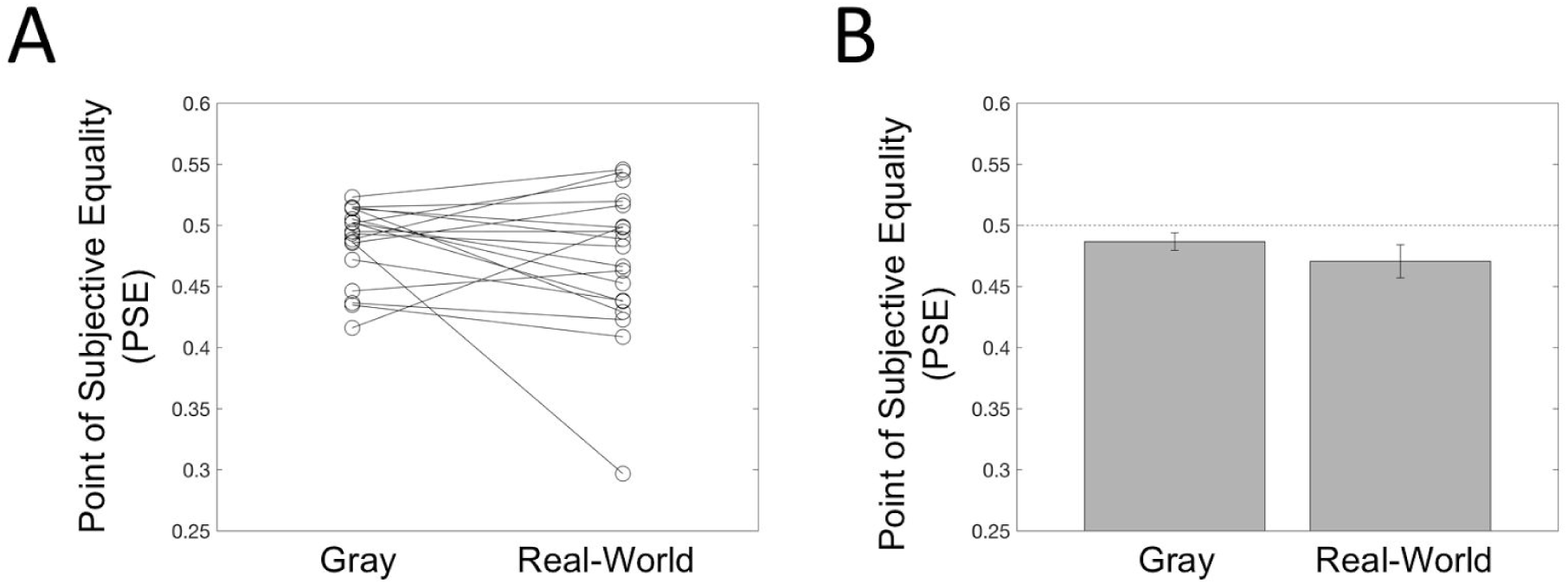
Estimates of the Point of Subjective Equality (PSE) for the Gray and Real-World conditions in Experiment 2. (A) Plotting PSEs for each individual subject for the Gray condition and Real-World condition. (B) Average PSEs for the two conditions. Bars display the mean PSE, and error bars represent the standard error of the mean of PSEs across all subjects. The PSEs in the Real-World condition (PSE mean: 0.4800) were significantly smaller than 0.5 (t(18) = −2.1638, p = 0.0442).

As in Experiment 1, we fitted psychometric curves to subjects’ data from each of two conditions, resulting in each subject having two psychometric fits. The criterion to discard subjects was the same as Experiment 1, with one additional element: if subjects reported that a background photo was not included in any of their trials (due to technical difficulties), or they did not fill in the post-experiment survey, data for both conditions for that participant were discarded. These standards resulted in four subjects’ data being excluded from further analysis (two subjects exhibited psychometric curves meeting the criterion for exclusion, one reported not seeing background photos during trials, and one chose not to fill out the post-experiment survey); we included the remaining 19 subjects in subsequent analyses. For the remaining subjects, as above we used the psychometric fits to calculate the PSE in each condition for each subject.

In contrast to our hypothesis that naturalistic backgrounds would amplify the subjective inflation of color saturation perception, results from a two-tailed, paired-sample, within-subjects t-test on the PSEs indicated that subjects perceived the peripheral stimuli similarly during the Gray condition compared to the Real-World condition similarly (t(18) =1.1643, p = 0.2595). We also conducted a 1-sample t-test of the PSE in each condition against 0.5 and found that the PSEs in the Gray condition (PSE mean: 0.4911) were not significantly different from 0.5, although there was a trend in the hypothesized direction (t(18) = −1.8703, p = 0.0778) akin to the results of Experiment 1. Finally, the PSEs in the Real-World condition (PSE mean: 0.4800) were significantly smaller than 0.5 (t(18) = −2.1638, p = 0.0442).

## Experiment 3

In considering what manipulations may lead to a larger effect size, we hypothesized that the magnitude of subjective inflation in the periphery may be directly proportional to the number of flashes. Therefore, we conducted a final experiment comparing 20 flashes to 1 flash, to test if this could also further increase the size of the effect observed in Experiment 1 where 10 flashes were compared to 1. Based on the results of Experiment 2 we also elected to present these stimuli with a background of naturalistic everyday images.

### Methods

#### Participants

Twenty-three participants were recruited online through Amazon Mechanical Turk. The payment protocol, consent procedure, exclusion criterion, and IRB approval procedures were identical to Experiments 1 and 2.

#### Stimuli & Procedure

Stimuli and procedure followed the same design used in Experiment 1, with a few differences. Instead of contrasting 1 flash versus 10 flashes, here we compared 1 flash versus 20 flashes to maximize the magnitude of the effect. We also employed fewer trials than Experiment 1, such that each participant completed 200 trials (2 conditions × 5 saturation levels × 20 trials per saturation level). As a result, the two one-minute breaks were given after trial 67 and after trial 133.

Due to the large amount of stimuli needed for this experiment, browser memory became a limiting factor. To circumvent this problem, we created 3 ‘pools’ of stimuli, with 30 images per saturation level in each set. Thus, in each trial, a random image from a random pool of the corresponding saturation level was chosen to be flashed (“1-flash” condition), or a random permutation of 20 images from the same pool were chosen and flashed (“20-flash” condition).

**Figure 7.**
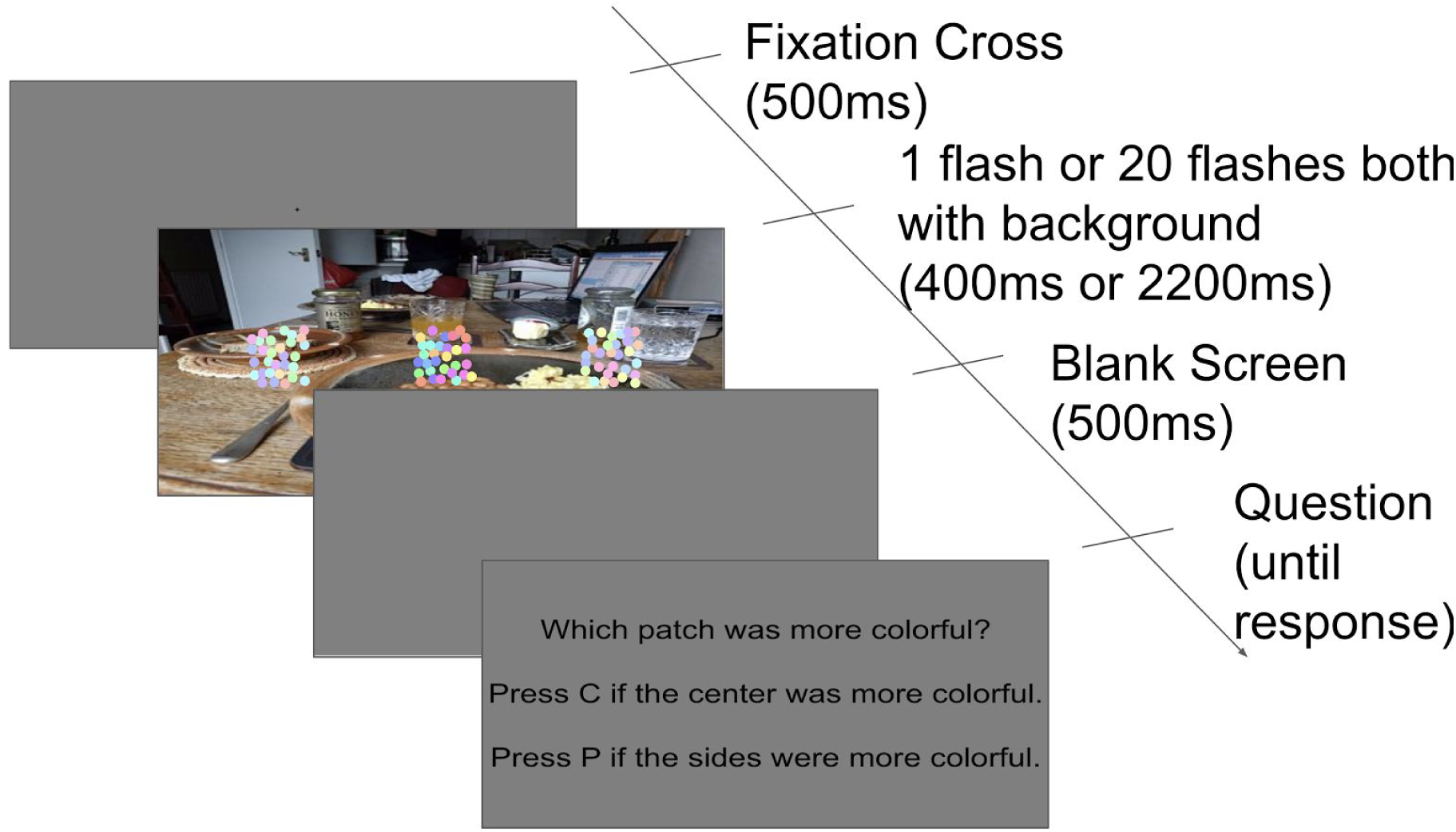
Stimulus presentation timeline in Experiment 3. The timeline in Experiment 3 was similar to Experiment 1, with the change that both conditions used an image of a real-world scene as background (akin to the Real-World condition in Experiment 2), and we used 20 flashes instead of 10 (“20-flash” condition).

#### Results

**Figure 8.**
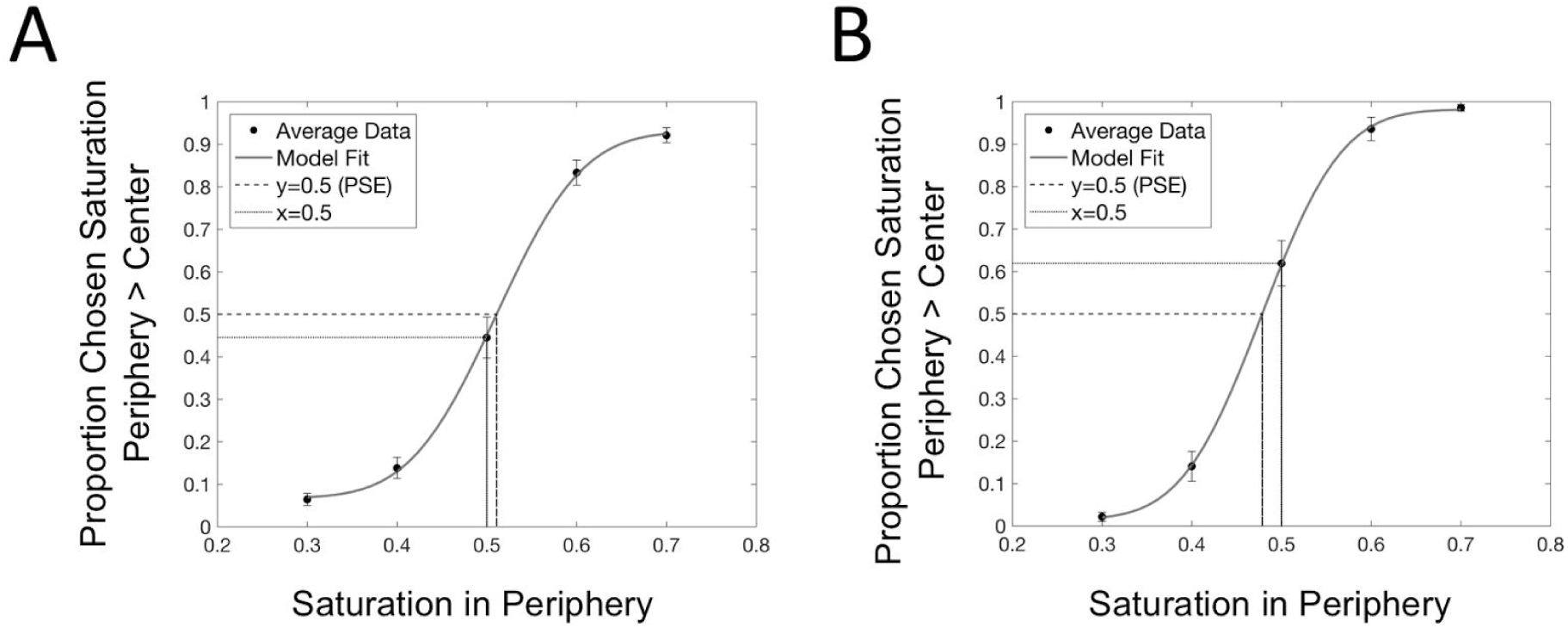
Psychometric fit to data averages across subjects. (A) Average psychometric fits for the 1-flash condition. Error bars represent standard error of the mean (SEM) across subjects. (B) Averaged psychometric fits for the 20-flash condition (right), with SEM across subjects. Results from our 2-tailed, paired-sample, within-subjects t-test on the Points of Subjective Equality (PSEs) indicate that subjects perceive the peripheral stimuli as being more saturated during the 20-flash condition compared to the 1-flash condition (t(20) =4.6875,p=0.00014).

**Figure 9.**
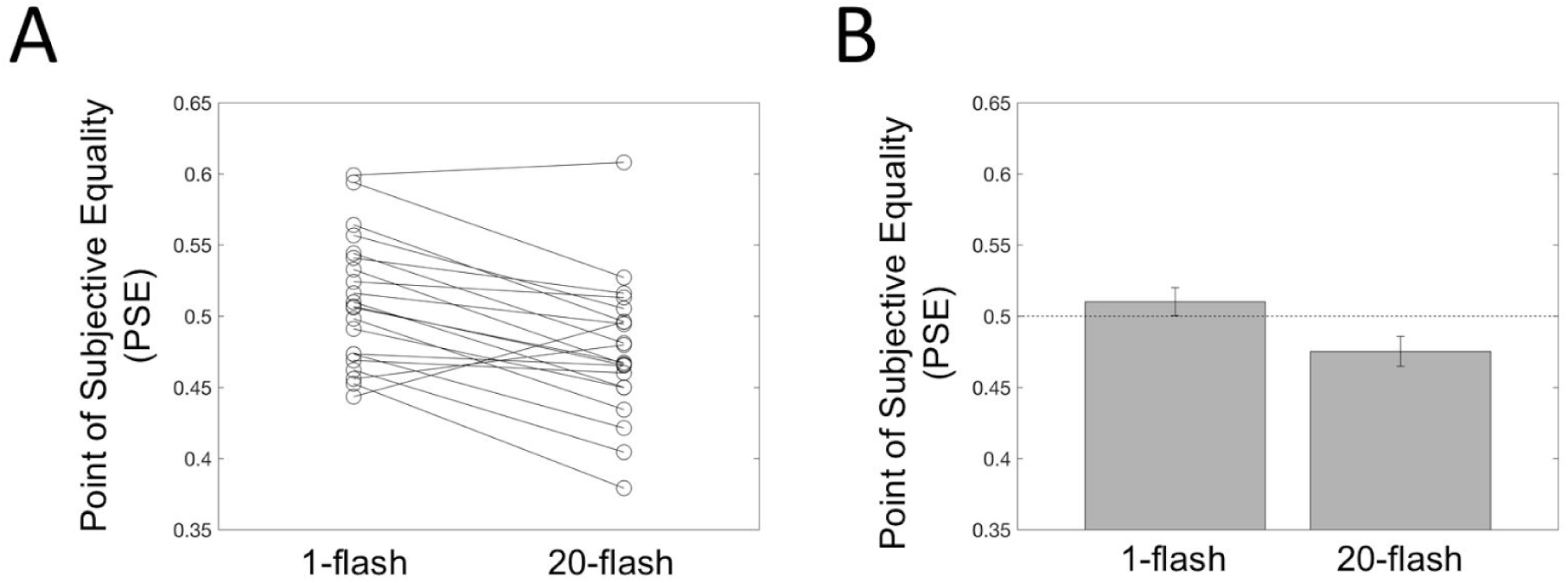
Estimates of the Point of Subjective Equality (PSE) for the 1-flash and 20-flash conditions. (A) PSEs for each individual subject for the 1-flash condition and 20-flash condition. (B) Average PSEs for the two conditions. Bars display the mean PSE, and error bars represent the standard error of the mean of PSEs across all subjects. We tested the PSEs of the two conditions against 0.5 with two 2-tailed 1-sample t-tests and found that the 1-flash condition (mean PSE: 0.5109) was not significantly different from 0.5 (t(20) = 1.0388, p = 0.3113) but the 20-flash condition (mean PSE: 0.4875) was significantly smaller than 0.5 (t(20) = −2.3503, p = 0.0291).

As in the previous experiments, we fit psychometric functions to the data of individual participants; curves that did not yield a meaningful Point of Subjective Equality (PSE) were discarded. If at least one of the two fitted psychometric curves of a participant has a range of < 0.2 or slope of <0.1, then the data of both conditions for that participant were discarded. This resulted in 2 subjects’ data being discarded, with the remaining 21 included in subsequent analyses. As above, for these subjects we used the psychometric fits to calculate the PSE for each condition for each subject.

Results from a 2-tailed, paired-sample, within-subjects t-test on the PSEs indicate that subjects perceive the peripheral stimuli as being more saturated during the 20-flash condition compared to the 1-flash condition (t(20) = 4.6875, p = 0.00014). We further tested the PSEs of the two conditions against 0.5 with two 2-tailed 1-sample t-tests and found that the 1-flash condition was not significantly different from 0.5 (mean PSE = 0.5109; t(20) = 1.0388, p = 0.3113) but the 20-flash condition was significantly smaller than 0.5 (mean PSE = 0.4875; t(20) = −2.3503, p = 0.0291).

## Discussion

Inspired by the dramatic and vivid nature of the flashed face distortion effect (Tangen et al. 2011), here we attempted to test whether a similar illusion can be created with simple color stimuli. We reasoned that there may be a common general mechanism for the two cases: under some degree of processing overload induced by repetitive flashes of stimuli, peripheral perception may start to be biased towards seeing things as more deviant from the background or norm. In the case of color, we tested if this means some simple patches of color dots may be perceived as more saturated. Conceptually, we confirmed our hypothesis (Experiment 1). In Experiment 2 we found that this effect was bigger under naturalistic background. In Experiment 3 we tried to see if we could obtain a bigger effect with higher number of flashes (20 flashes). While it appears that we could not obtain a bigger effect than would be obtained with natural background and 10 flashes only (Experiment 1 PSE = 0.4949 with 10 flashes and gray background, Experiment 2 PSE = 0.4800 with 10 flashes and naturalistic background, Experiment 3 PSE = 0.4875 with 20 flashes and naturalistic background), at the very least this shows that the effect is robust and can be replicated across experiments.

One potential concern about these experiments is that since they were conducted online, the experimental parameters were not controlled as much as would be possible in the lab. Thus subjects might be ‘cheating’ and looking at the side patches to maximize their performance. We acknowledge that while this is a valid concern, but we reason such cheating is unlikely because subjects had to focus on the central stimulus too, for the comparison, and that there are two peripheral stimuli (one on each side), meaning focusing one’s gaze on one would make seeing the other more difficult.

It could also be argued that the differences in the conditions are not phenomenological, but purely decisional. Ultimately, this is a psychophysical study and our measurements are forced-choice responses rather than open ended interviews concerning phenomenology. But at least this investigation is different from previous studies (Rahnev et al. 2011; Solovey et al. 2015) requiring subjects to detect the objective presence and absence of a stimulus. Here the responses concern the comparison of saturation. When saturation is illusorily perceived to be higher than it actually is, it is difficult to conceive of how this would not be reflected in phenomenological experience.

Finally, we note that the shifts in PSE here are quite small. We attempted in Experiment 2 and 3 to maximize this effect. While the shifts in PSE remain modest numerically, in terms of proportion of trials in which the peripheral stimuli were perceived as more saturated than the center one, the observed shifts are on the order of a bias of over 60%, which is perhaps not too insubstantial.

How does this compare to the vivid and dramatic nature of the original flashed face distortion effect? We note that for that original study, there were no quantitative data provided to facilitate a direct comparison. Introspectively, it seems like the effect in the face case should be bigger; perhaps this kind of effect works better for high level composite stimuli such as faces. One contribution of the present study is that we have introduced this PSE method for comparing the peripheral percept with a central stimulus, which can also be applied to the flashed face distortion effect (by asking subjects to compare where peripheral faces is more norm/caricature-like relative to a central standard). Future studies can capitalize on this method to allow more direct quantitative comparison between different stimulus domains, to allow us to understand the underlying mechanisms better.

Link to code and data:

https://github.com/vrsivananda/SubjectiveInflationInPeripheryExperiment

## References

Cohen, M. A., Dennett, D. C., & Kanwisher, N. (2016). What is the Bandwidth of Perceptual Experience? Trends in Cognitive Sciences, 20(5), 324–335. https://doi.org/10.1016/j.tics.2016.03.006

Dehaene, S., Changeux, J.-P., Naccache, L., Sackur, J., & Sergent, C. (2006). Conscious, *preconscious, and subliminal processing: a testable taxonomy*. Trends in Cognitive Sciences, 10(5), 204–211. https://doi.org/10.1016/j.tics.2006.03.007

Hansen, T., Pracejus, L., & Gegenfurtner, K. R. (2009). Color perception in the intermediate periphery of the visual field. Journal of Vision, 9(4), 26.1-12. https://doi.org/10.1167/9.4.26

Haun, A. M., Tononi, G., Koch, C., & Tsuchiya, N. (2017). Are we underestimating the richness of visual experience? Neuroscience of Consciousness, 3(1). https://doi.org/10.1093/nc/niw023

Kaunitz, L. N., Rowe, E. G., & Tsuchiya, N. (2016). Large Capacity of Conscious Access for Incidental Memories in Natural Scenes. Psychological Science, 27(9), 1266–1277. https://doi.org/10.1177/0956797616658869

Kouider, S., de Gardelle, V., Sackur, J., & Dupoux, E. (2010). How rich is consciousness? The partial awareness hypothesis. Trends in Cognitive Sciences, 14(7), 301–307. https://doi.org/10.1016/j.tics.2010.04.006

Lau, H., & Rosenthal, D. (2011). Empirical support for higher-order theories of conscious awareness. Trends in Cognitive Sciences, 15(8), 365–373. https://doi.org/10.1016/j.tics.2011.05.009

Peters, M. A. K., Ro, T., & Lau, H. (2016). Who’s afraid of response bias? Neuroscience of Consciousness, 2016(1). https://doi.org/10.1093/nc/niw001

Rahnev, D., Maniscalco, B., Graves, T., Huang, E., de Lange, F. P., & Lau, H. (2011). Attention induces conservative subjective biases in visual perception. Nature Neuroscience, 14(12), 1513–1515. https://doi.org/10.1038/nn.2948

Solovey, G., Graney, G. G., & Lau, H. (2015). A decisional account of subjective inflation of visual perception at the periphery. Attention, Perception & Psychophysics, 77(1), 258–271. https://doi.org/10.3758/s13414-014-0769-1

Tangen, J. M., Murphy, S. C., & Thompson, M. B. (2011). Flashed face distortion effect: grotesque faces from relative spaces. Perception, 40(5), 628–630. https://doi.org/10.1068/p6968

Vandenbroucke, A. R. E., Sligte, I. G., Barrett, A. B., Seth, A. K., Fahrenfort, J. J., & Lamme, V. A. F. (2014). Accurate metacognition for visual sensory memory representations. Psychological Science, 25(4), 861–873. https://doi.org/10.1177/0956797613516146

Witt, J. K., Taylor, J. E. T., Sugovic, M., & Wixted, J. T. (2015). Signal Detection Measures Cannot Distinguish Perceptual Biases from Response Biases. Perception, 44(3), 289–300. https://doi.org/10.1068/p7908

